# Neural signatures of value comparison in human cingulate cortex during decisions requiring an effort-reward trade-off

**DOI:** 10.1101/064105

**Authors:** Miriam C Klein-Flügge, Steven W Kennerley, Karl Friston, Sven Bestmann

## Abstract

Integrating costs and benefits is crucial for optimal decision-making. While much is known about decisions that involve outcome-related costs (e.g., delay, risk), many of our choices are attached to actions and require an evaluation of the associated motor costs. Yet how the brain incorporates motor costs into choices remains largely unclear. We used human functional magnetic resonance imaging during choices involving monetary reward and physical effort to identify brain regions that serve as a choice comparator for effort-reward trade-offs. By independently varying both options' effort and reward levels, we were able to identify the neural signature of a comparator mechanism. A network involving supplementary motor area (SMA) and the caudal portion of dorsal anterior cingulate cortex (dACC) encoded the difference in reward (positively) and effort levels (negatively) between chosen and unchosen choice options. We next modelled effort-discounted subjective values using a novel behavioural model. This revealed that the same network of regions involving dACC and SMA encoded the difference between the chosen and unchosen options' subjective values, and that activity was best described using a concave model of effort-discounting. In addition, this signal reflected how precisely value determined participants' choices. By contrast, separate signals in SMA and ventro-medial PFC (vmPFC) correlated with participants' tendency to avoid effort and seek reward, respectively. This suggests that the critical neural signature of decision-making for choices involving motor costs is found in human cingulate cortex and not vmPFC as typically reported for outcome-based choice. Furthermore, distinct frontal circuits ‘drive’ behaviour towards reward-maximization and effort-minimization.

**Significance Statement:** The neural processes that govern the trade-off between expected benefits and motor costs remain largely unknown. This is striking because energetic requirements play an integral role in our day-to-day choices and instrumental behaviour, and a diminished willingness to exert effort is a characteristic feature of a range of neurological disorders. We use a new behavioural characterization of how humans trade-off reward-maximization with effort-minimization to examine the neural signatures that underpin such choices, using BOLD MRI neuroimaging data. We find the critical neural signature of decision-making, a signal that reflects the comparison of value between choice options, in human cingulate cortex, whereas two distinct brain circuits ‘drive’ behaviour towards reward-maximization or effort-minimization.

## Introduction

Cost-benefit decisions are a central aspect of flexible goal-directed behaviour. One particularly well-studied neural system concerns choices where costs are tied to the reward outcomes (e.g., risk, delay; Kable and Glimcher, 2007; Boorman et al., 2009; Philiastides et al., 2010). Much less is known about choices tied to physical effort costs, despite their ubiquitous presence in human and animal behaviour. The intrinsic relationship between effort and action may engage neural circuits distinct from those involved in other value-based choice computations.

There is growing consensus that different types of value-guided decisions are underpinned by distinct neural systems, depending on the type of information that needs to be processed (e.g., Rudebeck et al., 2008; Camille et al., 2011b; Kennerley et al., 2011; Pastor-Bernier and Cisek, 2011; Rushworth et al., 2012). For example, activity in the ventro-medial prefrontal cortex (vmPFC) carries a signature of choice comparison (chosen-unchosen value) for decisions between abstract goods or when costs are tied to the outcome (Kable and Glimcher, 2007; Boorman et al., 2009; Fitzgerald et al., 2009; Philiastides et al., 2010; Hunt et al., 2012; Kolling et al., 2012; Clithero and Rangel, 2014; Strait et al., 2014). By contrast, such value difference signals are found more dorsally in medial frontal cortex when deciding between exploration versus exploitation (Kolling et al., 2012).

Choices requiring the evaluation of physical effort rest on representations of the required actions and their energetic costs, and thus likely require an evaluation of the internal state of the agent. This is distinct from choices based solely on reward outcomes (Rangel and Hare, 2010). Indeed, the proposed network for evaluating motor costs comprises brain regions involved in action planning and execution, including the cingulate cortex, putamen, and supplementary motor area (SMA) (Croxson et al., 2009; Kurniawan et al., 2010; Prévost et al., 2010; Kurniawan et al., 2013; Burke et al., 2013; Bonnelle et al., 2016). Neurons in anterior cingulate cortex (ACC) encode information about rewards, effort costs, and actions (Matsumoto et al., 2003; Kennerley and Wallis, 2009; Luk and Wallis, 2009; Hayden and Platt, 2010), and integrate this information into an economic value signal (Hillman and Bilkey, 2010; Hosokawa et al., 2013). Moreover, lesions to ACC profoundly impair choices of effortful options and between action values (Walton et al., 2003, 2006, 2009; Schweimer and Hauber, 2005; Kennerley et al., 2006; Rudebeck et al., 2006, 2008; Camille et al., 2011b).

While these studies highlight the importance of motor-related structures in representing effort information, it remains unclear whether computations in these regions are indeed related to *comparing* effort values (or effort-discounted net values) – the essential neural signature which would implicate these areas in decision making. Indeed, these regions could simply *represent* effort which is then passed onto other regions for value comparison processes. A number of questions thus arise. First, is information about reward and effort compared in separate neural structures, or is this information fed to a region that compares options based on their integrated value? Second, do regions that preferably encode reward or effort have a direct influence on determining choice? Finally, assuming separate neural systems are present for influencing choices based on reward versus effort, how does the brain arbitrate between these signals when reward and effort information support opposing choices?

Here we employed a task designed to identify signatures of a choice comparison for effort-based decisions in humans using fMRI, and to test whether different neural circuits ‘drive’ choices towards reward-maximization versus energy-minimization. We show that the neural substrates of effort-based choice are distinct from those computing outcome-related choices: well-known reward and effort circuits centred on vmPFC and SMA bias choices to be more driven by benefits or motor costs, respectively, with a region in cingulate cortex integrating cost and benefit information and comparing options based on these integrated subjective values.

## Materials and methods

### Participants

24 participants with no history of psychiatric or neurological disease, and with normal or corrected-to-normal vision took part in this study (mean age 28±1, age range 19-38, 11 females). All participants gave written informed consent and consent to publish prior to the start of the experiment; the study was approved by the local research ethics committee at University College London (UCL; 1825/003) and conducted in accordance with the declaration of Helsinki. Participants were reimbursed with £15 for their time and in addition they accumulated average winnings of £7.16±0.11 during each of the two blocks of the task (the maximum winnings per block were scaled to £8; the resulting average total pay was £29.32). Three participants were excluded from the analysis, one for failing to stay awake during scanning, and two due to excessive head movements (summed movement in any direction and run >40mm). All analyses were performed on the remaining 21 participants.

### Behavioural task

Participants received both written and oral task instructions. They were asked to make a series of choices between two options, which independently varied in required grip force (‘effort’) and reward magnitude (**Figure 1A**). The reward magnitude was shown as a number (range: 10-40 points; roughly corresponding to pence) and required force levels were indicated as the height of a horizontal bar (range: 20-80% of the participant's maximum grip force).

**Figure 1.**
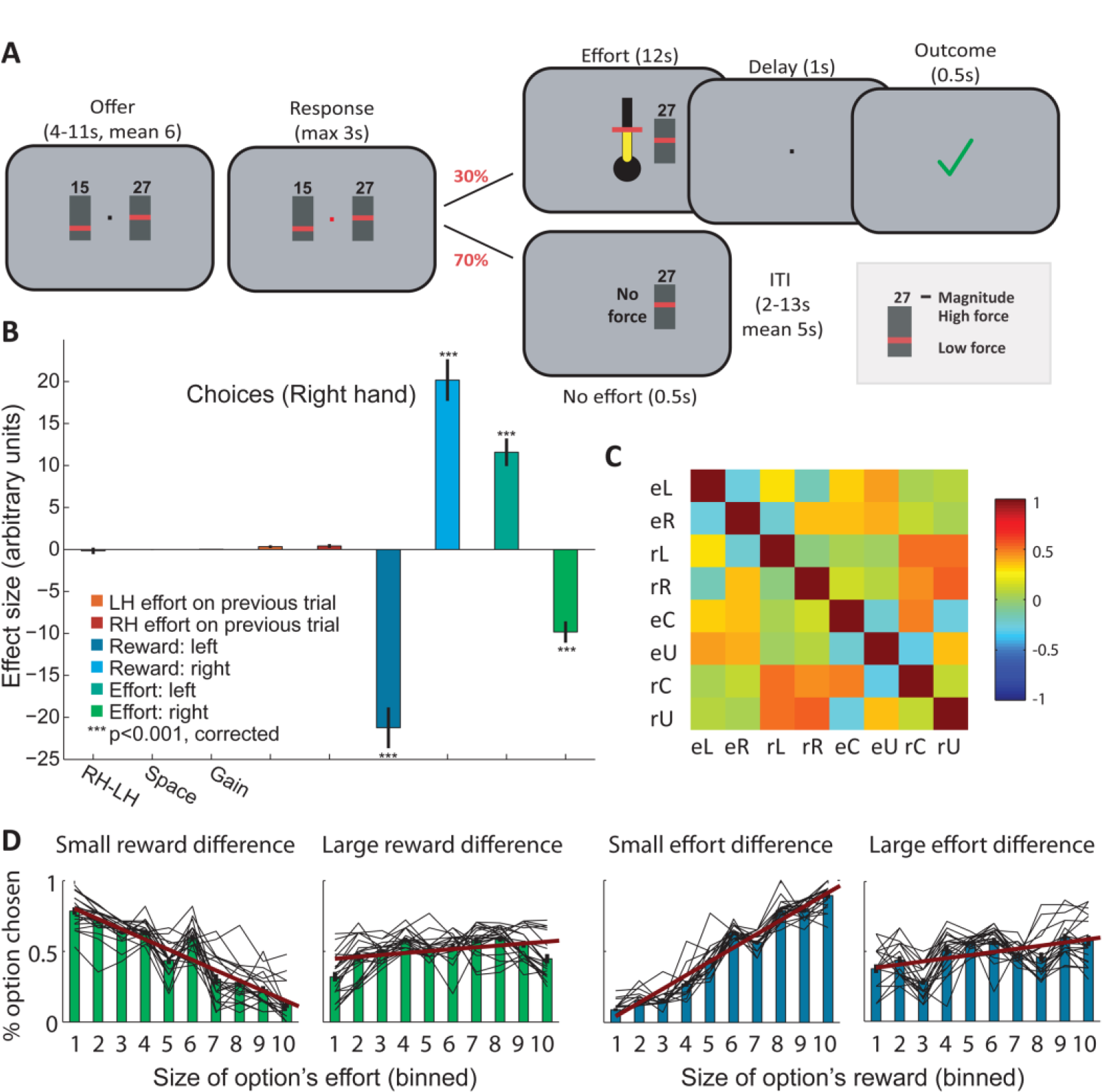
Task and Behaviour. **A**, Human participants chose between two options associated with varying reward magnitude (numbers) and physical effort (bar height translates intoforce,‘Offer’). Once the fixation cross turned red (‘Response’), participants were allowed to indicate their choice. Thus, the time of choice computation was separable in time from the motor response. Following aresponse,the effort had to be realised on an unpredictable 30% of trials (top). On thesetrials,participants had to produce a 12s power grip at a strength proportional to the bar height of the chosen option. Force levels were adjusted to individuals' maximum force at the start of the experiment. Participants received feedback about successful performance of the grip (99% accuracy), and the rewards collected on successful trials were added to the total winnings. On 70% of trials (bottom), no effort was required and the next trial commenced (Inter-trial interval; ITI). **B**, Participants' choices were driven by both options' reward magnitude and effort level showing that all dimensions of the outcome were taken into account for computing a choice. Benefits and costs had opposite effects: larger efforts discouraged and larger reward magnitudes encouraged the choice of an option. Standard errors denote ±SEM.**C**, Correlations between left (L), right (R), chosen (C), and unchosen (U) effort levels (e) and reward magnitudes (m) show that the regressors of interest were sufficiently decorrelated in our design. **D**, Effort has a strong effect on choice in trials with small rewarddifferences,but no effect when the reward difference is large (green panels; median split on reward difference; effort binned for visualization). Similarly, reward has a stronger effect in trials with small effort differences compared to trials with large effort differences (blue panels). This shows that participants indeed trade-off effort against reward and confirms that reward has a stronger and opposite effect compared to effort (red slope), as shown in **B**. The black lines correspond to individual participants and suggest that reward and effort were treated as continuous variables.

Each trial comprised an offer, response and outcome phase; a subset of 30% of trials also contained an effort production phase. During the ‘offer’ phase, participants decided which option to choose but they were not yet able to indicate their response. There were two trial types (50% each): ‘ACT’ (action) and ‘ABS’ (abstract). In ACT trials, the two choice options were presented to the left and right of fixation, and thus in a horizontal or ‘action space’ configuration in which the side of presentation directly related to the hand with which to choose that option. In ABS trials, choice options were shown above and below fixation, and thus in a vertical or ‘goods space’ arrangement that did not reveal the required action. In both conditions, stimuli were presented close to the centre of the screen and participants did not need to move their eyes to inspect them. To maximally distinguish the hemodynamic response from the offer and response phase, the duration of the offer phase varied between 4-11s (Poisson distributed; mean 6s).

The response phase started when the fixation cross turned red. In ACT trials, the arrangement of the two choice options remained the same; in ‘ABS’ trials, the two options at the top and bottom were switched to the left and right of fixation (with a 50/50% chance), thus revealing the required action mapping. Choices were indicated by a brief squeeze of a grip device (see below for details) on the corresponding side (maximum response time: 3 s; required force level: 35% of maximum voluntary contraction; MVC). Note that ACT and ABS trials were merged for all analyses because no significant differences were found for the tests reported in this manuscript.

On 70% of trials, no effort was required: as soon as participants indicated their choice, the unchosen option disappeared, and the message ‘no force’ was displayed for 500ms. The next trial commenced after a variable delay (ITI: 2-13s; Poisson distributed; mean: 5s). On the remaining 30% of trials, a power grip of 12s was required (‘effort’). Again, the unchosen option disappeared but now a thermometer appeared centrally and displayed the target force level of the chosen option. Participants were given on-line visual feedback about the applied force level using changing fluid levels in the thermometer. On successful application of the required force for at least 80% of the 12s period, a green tick appeared (500ms; ‘outcome’ phase; delay preceding outcome: 0.5-1.5s uniform) and the reward magnitude of the chosen option was added to the total winnings. Otherwise, the total winnings remained unchanged (red cross: 500ms). Because participants were almost always successful in applying the required force on effort trials (accuracy: 99.30α0.004%; only four participants made any mistakes), there was no confound between effort level and risk/reward expectation.

The sensitivity of the grip device was manipulated between trials (‘high’ or ‘low’). A high gain meant that the grippers were twice as sensitive as for a low gain, and thus the same force deviation doubled the rate of change in the thermometer's fluid level. While this manipulation was introduced to study interactions between mental and physical effort, none of our behavioural or fMRI analyses revealed any significant effects of gain during the choice phase, which is the focus of the present paper.

To summarize, our task involved several important features: (a) as our aim was to specifically examine value comparison mechanisms during effort-based choice, we manipulated both options' values and thus the expected values of the two offers had to be computed and compared on-line in each trial, unlike in previous experiments (Croxson et al., 2009; Kurniawan et al., 2010, 2013; Prévost et al., 2010; Burke et al., 2013; Bonnelle et al., 2016); (b) the decision process and the resulting motor response were separated in time (**Figure 1A**). This enabled us to examine the value comparison in the absence of-and not confounded with-processes related to action execution; (c) both reward and effort levels were varied parametrically rather than in discrete steps, and orthogonally to each other, thereby granting high sensitivity for the identification of effort and reward signals, respectively; (d) efforts were only realised on a subset of trials, ensuring that decisions were not influenced by fatigue (Klein-Flügge et al., 2015). Importantly, however, at the time of choice participants did not know whether a given trial was real or hypothetical and therefore the optimal strategy was to treat each trial as potentially real; (e) the duration of the grip on effort trials (12s) had been determined in pilot experiments and ensured that force levels were factored into the choice process. Moreover, the fixed duration of grip force also meant that effort costs were not confounded with temporal costs.

### Scanning procedure

Prior to scanning, force levels were adjusted to each individual's grip strength using a grip calibration. Participants were seated in front of a computer monitor and held a custom-made grip device in both hands. Each participant's baseline (no grip) and MVC were measured over a period of 3 s, separately for both hands. The measured values were used to define individual force ranges (0-100%) for each hand, which were then used in the behavioural task, both pre-scanning and during scanning.

Before entering the scanner, participants completed a training session consisting of one block of the behavioural task (112 trials, ~30 minutes). This gave them the opportunity to experience different force levels and to become familiar with the task. Importantly, it also ensured that decisions made subsequently in the scanner would not be influenced by uncertainty about the difficulty of the displayed force levels. In the scanner, participants completed two blocks of the task (overall task duration ~60 minutes; 224 choices).

### Generation of choice stimuli

Because our main question related to the encoding of value difference signals during effort-based choices, the generation of suitable choice stimuli was a key part of the experimental design. Choice options were identical for every individual and were chosen such that they would minimise the correlation between the fMRI regressors for chosen and unchosen effort, reward magnitude and value (obtained mean correlations post-scanning: effort: −0.23; reward magnitude: 0.11; value: 0.43; **Figure 1C**). We also ensured that left and right efforts, reward magnitudes and values were decorrelated to be able to identify action value signals (effort: 0.28; reward magnitude: 0.05; value: 0.07). We simulated several individuals using a previously suggested value function for effort-based choice (Prévost et al., 2010). Stimuli were optimized with the following additional constraints: either the efforts or the reward magnitudes had to differ by at least 0.1 on each trial, the range of efforts and reward magnitudes was [0.2 to 0.8] × MVC or 0-50 points, respectively, and the overall expected value for both hands was comparable. Furthermore, in 85% of trials the larger reward was paired with the larger effort level, and the smaller reward with the smaller effort level, making the choice hard, but on 15% of trials, the larger reward was associated with the smaller effort level (‘no-brainer’). The two choice sets that minimized the correlations between our regressors of interest were used for the fMRI experiment. A third stimulus set was saved for the behavioural training prior to scanning.

Preliminary fMRI analyses revealed that we had overlooked a bias in our stimuli. In the last third of trials of the second block, the overall offer value ((magnitude1/effort1 + magnitude2/effort 2)/2) decreased steadily, leading to skewed contrast estimates. Therefore the last 40 trials were discarded from all analyses.

Note that we refer to choices in this study as ‘effort-based’ to highlight the distinction from purely outcome/reward-based choices or choices involving other types of costs (e.g., delay-based). But of course, in our task, all choices were effort-as well as reward-based.

### Recordings of grip strength

The grippers were custom-made and consisted of two force transducers (FSG15N1A, Honeywell, NJ, USA) placed between two moulded plastic bars (see also Ward and Frackowiak, 2003). A continuous recording of the differential voltage signal, proportional to the exerted force, was acquired, fed into a signal conditioner (CED 1902, Cambridge Electronic Design, Cambridge, UK), digitized (CED 1401, Cambridge Electronic Design, Cambridge, UK) and fed into the computer running the stimulus presentation. This enabled us, during effort trials, to give online feedback about the exerted force using the thermometer display.

### Behavioural analysis

To examine which task variables affected participants' choice behaviour, a logistic regression was fitted to participants' choices (1=RH, 0=LH) using the following nine regressors: a RH-LH bias (constant term); condition (ABS or ACT); gain (high or low); LH-effort on previous trial; RH-effort on previous trial; reward magnitude left; reward magnitude right; effort left; effort right. T-tests performed across participants on the obtained regression coefficients were adjusted for multiple comparisons using Bonferroni correction. Since only reward magnitudes and efforts influenced behaviour significantly (see Results), the logistic regression models performed for the analysis of the neural data below (equations (2) and (3)) only contained these variables (or their amalgamation into combined value).

To examine the influence of reward and effort on participants' choice behaviour in more depth, we tested whether participants indeed weighed up effort against reward, and whether they treated reward and effort as continuous variables. If reward and effort compete for their influence on choice, then the influence of effort should become larger as the reward difference becomes smaller, and vice versa. Thus, we performed a median split of our trials according to the absolute difference in reward (or effort) between the two choice options. We then calculated the likelihood of choosing an option as a function of its effort (reward) level, separately for the two sets of trials. Effort (reward) values were distributed across ten bins with equal spacing; this binning was independent of the effort (reward) level of the alternative option. For statistical comparisons, we fitted a slope for each participant to the mean of all bins. T-tests were performed on the resulting four slopes testing for the influence of (a) effort in trials with small reward difference, (b) effort in trials with large reward difference, (c) reward in trials with small effort difference, and (d) reward in trials with large effort difference. We report uncorrected p-values but all conclusions hold when correcting for six comparisons (a-d against zero; a versus b; c versus d).

We also tested for effects of fatigue: the above logistic regression suggested that choices were not affected by whether or not the previous trial required the production of effort, as shown previously in this task (Klein-Flügge et al., 2015). More detailed analyses examined the percentage of trials in which the higher effort option was chosen (running average across 20 trials), and participants' performance in reaching and maintaining the required force. The latter was measured as the time point when 10 consecutive samples were above force criterion (the shorter, the sooner), and as the percentage of time out of 12s that participants were at criterion, respectively. For all measures, we compared the first and last third of trials. Here we report the comparison between the first and last third across the entire experiment. However, separate analyses-using the first and last third of just the first or the second block-revealed identical results. There were no effects of fatigue: in all cases participants either improved or stayed unchanged (% higher effort chosen: first third 60.56%±1.94; last third 60.93%±2.79, p=0.69, t_20_=-0.40; reaching the force threshold: first third 0.83s±0.04; last third 0.76s±0.03; p=0.01, t_20_=2.82; maintaining the force above threshold: first third 92.49%±0.47; last third 93.51%±0.28; p=0.01, t_20_=-2.84).

To derive participants' subjective values for the offers presented on each trial, we developed an effort discounting model (Klein-Flügge et al., 2015). This model has been shown to provide better fits than the hyperbolic model previously suggested for effort discounting (Prévost et al., 2010) both here and in our previously published work (Klein-Flügge et al., 2015). Crucially, its shape is initially concave, unlike a hyperbolic function, allowing for smaller devaluations of value for effort increases at weak force levels, and steeper devaluations at higher force levels, which is intuitive for effort discounting and biologically plausible. Our model follows the following form:

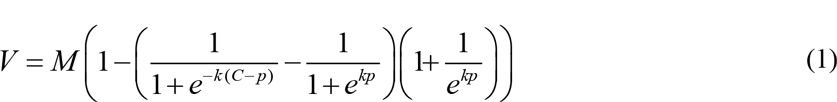

*V* is subjective value, *C* is the effort cost, *M* the reward magnitude, and *k* and *p* are free parameters. *C* and *M* are scaled between 0 and 1, corresponding to 0% MVC and 100% MVC, and 0 points and 50 points, respectively. A simple logistic regression on the difference in subjective values between choice options was then used to fit participants' choices; in other words, the following function (‘softmax rule2019) was used to transform the subjective values *V1* and *V2* of the two options offered on each trial into the probability of choosing option 1.

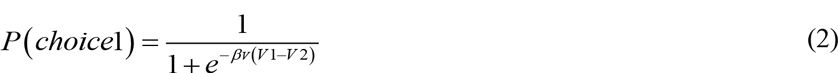

The free parameters (slope *k*, turning point *p*, softmax precision parameter *β_v_*), were fitted using the Variational Laplace algorithm (Penny et al., 2003; Friston et al., 2007). This is a Bayesian estimation method which incorporates Gaussian priors over model parameters and uses a Gaussian approximation to the posterior density. The parameters of the posterior are iteratively updated using an adaptive step size, gradient ascent approach. Importantly, the algorithm also provides the free energy *F*, which is an approximation to the model evidence. The model evidence is the probability of obtaining the observed choice data, given the model. To maximize our chances to find global, rather than local maxima with this gradient ascent algorithm, parameter estimation was repeated over a grid of initialization values, with eight initializations per parameter. The optimal set of parameters, i.e., that obtained from the initialization that resulted in the maximal free energy, was used for modelling subjective values in the fMRI data.

For our BOLD analyses, the only relevant parameter was *β_v_*. It reflects the weight (i.e., strength) with which participants' choices are driven by subjective value, rather than noise; it is also often referred to as precision or inverse softmax temperature. Fitting ACT and ABS, or high and low gain trials separately did not lead to any significant differences between conditions (paired t-tests on parameter estimates between conditions all p>0.3), and did not improve the model evidence (paired t-test on the model evidence; fitting conditions separately or not: p=0.82; fitting gain separately or not: p=0.63). Trials were therefore pooled for model fitting. Once fitted, the performance of our new model was compared to that of the hyperbolic model and two parameter-free models (difference: reward-effort; quotient: reward/effort) as described in (Klein-Flügge et al., 2015) using a formal model comparison.

### FMRI data acquisition and pre-processing

The fMRI methods followed standard procedures (e.g., Klein-Flügge et al., 2013): T2*-weighted echo-planar images (EPI) with blood oxygenation level-dependent (BOLD) contrast were acquired using a 12-channel head coil on a 3Tesla Trio MRI scanner (Siemens, Erlangen, Germany). A special sequence was used to minimise signal drop out in the OFC region (Weiskopf et al., 2006) and included an echo time (TE) of 30ms, a tilt of 30° relative to the rostro-caudal axis and a local z-shim with a moment of −0.4 mT/m ms applied to the OFC region. To achieve whole-brain coverage, we used 45 transverse slices of 2mm thickness, with an interslice gap of 1mm and in-plane resolution of 3×3 mm, and collected slices in an ascending order. This led to a repetition time (TR) of 3.15 seconds. In each session, a maximum of 630 volumes were collected (~33 minutes) and the first five volumes of each block were discarded to allow for T1 equilibration effects. A single T1-weighted structural image with 1mm^3^ voxel resolution was acquired and co-registered with the EPI images to permit anatomical localisation. A fieldmap with dual echo-time images (TE1= 10ms, TE2 = 14.76ms, whole brain coverage, voxel size 3×3×3 mm) was obtained for each subject to allow for corrections in geometric distortions induced in the EPIs at high field strength (Andersson et al., 2001).

During the EPI acquisition, we also obtained several physiological measures. The cardiac pulse was recorded using an MRI-compatible pulse oximeter (Model 8600 F0, Nonin Medical, Inc. Plymouth, MN, USA), and thoracic movement was monitored using a custom-made pneumatic belt positioned around the abdomen. The pneumatic pressure changes were converted into an analogue voltage using a pressure transducer (Honeywell International Inc. Morristown, NJ, USAir) before digitization, as reported in (Hutton et al., 2011).

Pre-processing and statistical analyses were carried out using SPM8 (Wellcome Trust Centre for Neuroimaging, London, UK, www.fil.ion.ucl.ac.uk/spm). Image pre-processing consisted of realignment of images to the first volume, distortion correction using fieldmaps, slice time correction, conservative independent component analysis to identify and remove obvious artefacts (using MELODIC in Fmrib's Software Library, http://fsl.fmrib.ox.ac.uk/), coregistration with the structural scan, normalisation to a standard MNI template, and smoothing using an 8mm full-width at half maximum Gaussian kernel.

### Data analysis – General Linear Model

The first general linear model (GLM1) included twelve main event regressors. The offer phase was described using onsets for (i) ACT trials preparing a left response; (ii) ACT trials preparing a right response; and (iii) ABS trials. All three events were modelled using durations of 2s and were each associated with four parametric modulators: the reward magnitude and effort of the chosen and unchosen option. Crucially, these four parametric modulators competed to explain common variance during the estimation, rather than being serially orthogonalised (in other words, we implicitly tested for effects that were unique to each parametric explanatory variable). The response phase was described using four regressors for ‘no force’ trials (1s duration) and four regressors for effort production trials (12s duration): (iv-vii) no force ACT_left_, ACT_right_, ABS_Left_ ABS_right_; (viii-xi) effort production left (low gain), left (high gain), right (low gain), right (high gain). Finally, the outcome was modelled as a single regressor because the proportion of trials in which efforts were not produced successfully was negligible (median: 0; mean: 0.43 ± 0.22 trials; only 4 of 21 participants had any unsuccessful trials).

In addition to event regressors, a total of 23 nuisance regressors were included to control for motion and physiological effects of no interest. First, to account for motion-related artefacts that had not been eliminated in rigid-body motion correction, the six motion regressors obtained during realignment were included. Second, to remove variance accounted for by cardiac and respiratory responses, a physiological noise model was constructed using an in-house Matlab toolbox (Hutton et al., 2011). Models for cardiac and respiratory phase and their aliased harmonics were based on RETROICOR (Glover et al., 2000). The model for changes in respiratory volume was based on (Birn et al., 2006). This resulted in 17 physiological regressors in total: ten for cardiac phase, six for respiratory phase, and one for respiratory volume.

The parameters of the hemodynamic response function (HRF) were modified to obtain a double-gamma HRF, with the standard settings in Fmrib's Software Library (http://fsl.fmrib.ox.ac.uk/): delay to response 6, delay to undershoot: 16, dispersion of response: 2.5; dispersion of undershoot: 4; ratio of response to undershoot: 6, length of kernel: 32 (all in seconds).

The second GLM (GLM2) was identical to the first, except that the four parametric regressors (reward magnitude and effort of the chosen and unchosen option) were replaced by the subjective model-derived values of the chosen and unchosen option. This allowed us to identify regions encoding the difference in subjective value between the offers.

Three further GLMs were fitted to the data to test whether the values derived from the sigmoidal model provide the best explanation of the measured BOLD signals. These GLMs were identical to GLM2 except that the parametric regressors for the values of the chosen and unchosen option derived from the sigmoidal model were replaced by either (a) the values derived from a hyperbolic model (GLM3), (b) the values derived from a parameter-free difference ‘reward-effort’ (GLM4), or (c) the values derived from a parameter-free quotient ‘reward/effort’ (GLM5).

### Identifying signatures of choice computation

Our first aim was to identify brain regions with BOLD signatures of choice computation (**Figure 2A**). Thus, we first identified brain regions that fulfilled the following two criteria (GLM1): (a) the BOLD signal correlated negatively with the difference in effort between chosen and unchosen options, and (b) the BOLD signal correlated positively with the difference in reward magnitude between chosen and unchosen options. Collectively, these two signals form the basis of a value difference signal because effort contributes negatively and reward magnitude contributes positively to overall value. Previous work has demonstrated, using predictions derived from a biophysical cortical attractor network, that at the level of large neural populations, as measured using human neuroimaging techniques such as fMRI or MEG, the characteristic signature of a choice comparison process is a value difference signal (Hunt et al., 2012). The responses predicted for harder and easier choices differ because the speed of the network computations varies as a function of choice difficulty (e.g., faster for high value difference). Thus, an area at the formal conjunction of the two contrasts described by (a) and (b) would carry the relevant signatures for computing a subjective value difference signal, a cardinal requirement for guiding choice. Importantly, while we reasoned that the choice computations in our specific task should follow similar principles as in Hunt et al. (2012), we expected this computation to occur in different regions because it would be based on the integration of a different type of decision cost. In an additional analysis (**Figure 5**), for completeness, we also identified brain regions significant in the inverse contrast, i.e., a conjunction of positive effort and negative reward magnitude difference (Wunderlich et al., 2009; Hare et al., 2011).

**Figure 2.**
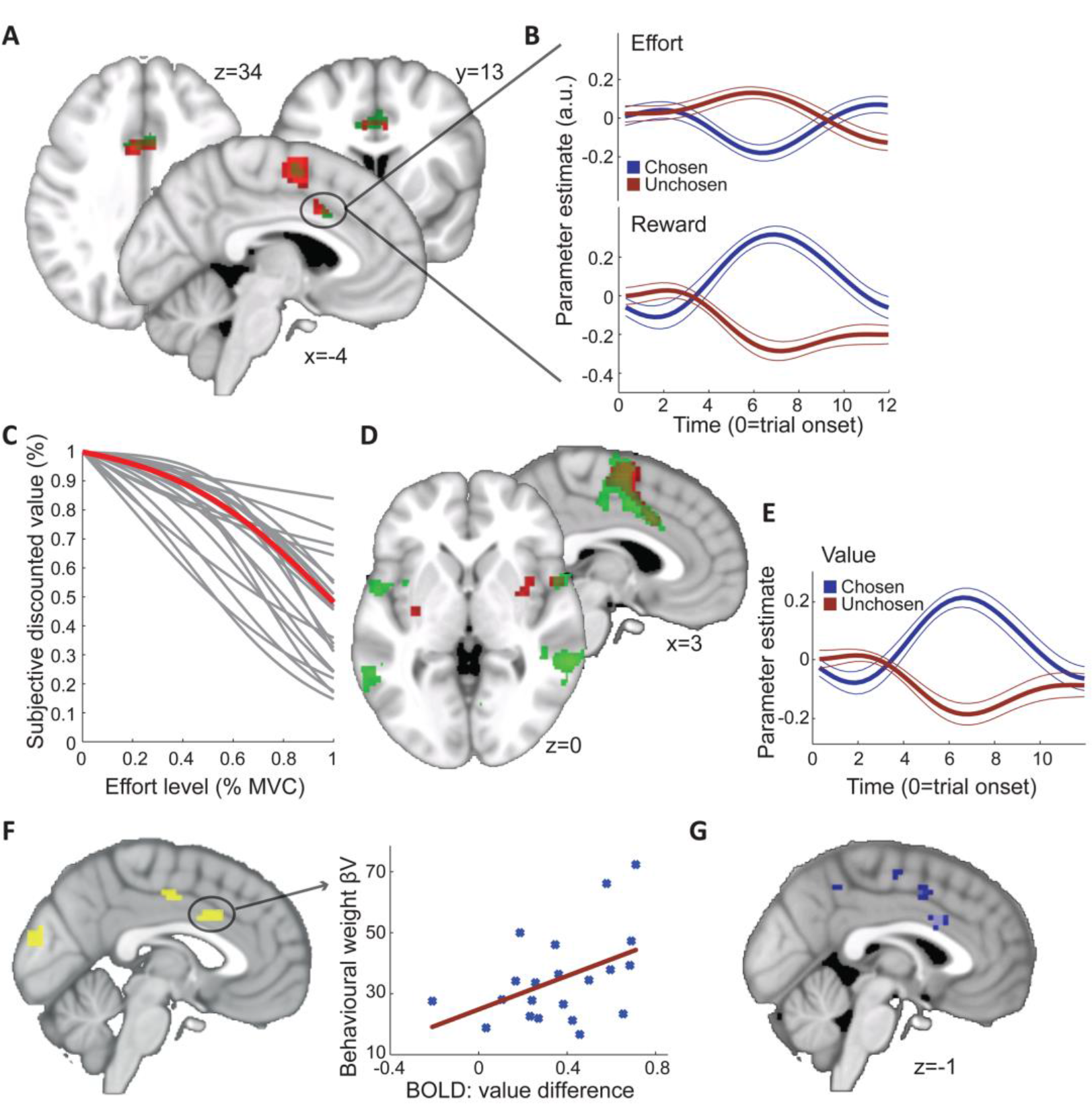
Neural signatures of effort choice comparison in SMA and dACC. **A**, As a marker of choice computation,we identified regions encoding (a) the difference between the chosen and unchosen reward magnitudes and (b) the inverse difference between the chosen and unchosen effort levels. The conjunction of both contrasts in SPM (shown at p<0.001 uncorr.) revealed the supplementary motor area (SMA) and a region in the caudal portion of dorsal anterior cingulate cortex (dACC) (both survive FWE-corr p<0.05). Cluster-level corrected results obtained from FSL's Flame 1 (z>2.3, p<0.05) are overlaid in green to confirm this finding. **B**, For illustration purposes,the two opposing difference signals are shown for the dACC cluster on the right. Standard errors denote ± SEM. **C**, A custom-built sigmoidal model was fitted to participants' choices to obtain individual effort discounting curves (grey; red: group mean). In the model,the subjective value of an option's reward (y-axis, represented in %) is discounted with increasing effort levels (x-axis). This allowed inferring the subjective values ascribed to choice options and modelling of subjective value in the BOLD data. **D**, The difference in subjective value between the chosen and unchosen option,as derived from the behavioural effort discounting model inC,was encoded in a similar network of regions as the combined difference in reward magnitude and effort shown inA,including caudaldACC,SMA, bilateral putamen and insula (shown at p<0.001 uncorr. as obtained with SPM; cluster-level corrected FSL results overlaid in green for z>2.7, p<0.05). **E**, The subjective value difference signal extracted from the dACC is shown for illustration (standard errors: ± SEM). **F**, Left: Regions encoding subjective value as in D but where the strength of this signal additionally correlated with the extent to which value difference guided behaviour (inverse softmax temperature β_v_; shown at p<%0.01 uncorr.; only the dACC survives cluster-level FWE-corr p<0.05). Right: Illustration of the correlation in dACC for visual display purposes only. The stronger the BOLD difference between the chosen and unchosen option in thisregion,the more precisely participants' choices are guided by value (β_v_). This suggests that the dACC's value signal computed at the time of choice is relevant for guiding choices. **G**, Regions where the encoding of effort differencecorrelates, acrosssubjects,with a marker for the individual level of ‘effort distortion’ as captured by the parameters *k* and *p* of the modelled discount function. The better an individual's subjectively experienced effort was captured in the GLM (i.e., the less distorted their discount function), the stronger the inverse effort difference signal in caudal dACC and SMA (light blue: p<0.001 uncorr; dark blue: p<0.005 uncorr; dACC /SMA survive cluster-level FWE-corr p<0.05). This suggests dACC and SMA encode effort difference in the way it subjectively influences the choice.

### Regions of Interest and extraction of time courses

For whole-brain analyses we used a FWE cluster-corrected threshold of p<.05 (using a cluster-defining threshold of p<0.01 and a cluster threshold of 10 voxels). For a priori region of interest (ROI) analyses we used a small-volume corrected FWE cluster-level threshold of p<0.05 in spheres of 5mm around previous coordinates, namely in left and right putamen ([±26, −8, −2] (Croxson et al., 2009)), SMA ([4 −6 58], (Croxson et al., 2009)) and vmPFC ([−6, 48, −8] (Boorman et al., 2009)).

BOLD time series were extracted from the pre-processed data of the identified regions by averaging the time series of all voxels that were significant at p<0.001 (uncorrected). Time series were up-sampled with a resolution of 315ms (1/10*TR) and split into trials for visual illustration of the described effects (e.g., **Figure 2B**).

At the suggestion of one reviewer, the two main analyses (conjunction of reward and inverse effort difference described above, and value difference contrast described below) were repeated in FSL using Flame1 because of differences between SPM and FSL in controlling for false positives when using cluster-level corrections (Eklund et al., 2015). For this control analysis, we imported the pre-processed (unsmoothed) images to FSL. We then used FSL's default smoothing kernel of 5mm and a cluster-forming threshold of z>2.3 (corresponding to p<0.01; default in FSL). The obtained results are overlaid in **Figure 2A** and **2D**.

### Encoding of subjective value

We next asked whether BOLD signal changes in the regions identified using the abovementioned conjunction could indeed be described by the subjective values derived from our custom-made behavioural model. We thus performed a whole-brain contrast identifying regions encoding the difference in subjective value between the chosen and unchosen option (GLM2; **Figure 2D**). To test whether the BOLD signal was better explained by subjective value as modelled using the sigmoidal function or three alternative models (hyperbolic; ‘difference’: reward-effort; ‘quotient’: reward/effort; GLM3-GLM5; **Figure 3B**), we calculated the difference between the value difference maps obtained on the first level for each participant (sigmoid vs hyperbolic; sigmoid vs difference; sigmoid vs quotient; **Figure 3C**). A standard second-level t-test was performed on the three resulting difference images and statistical significance evaluated as usual.

**Figure 3.**
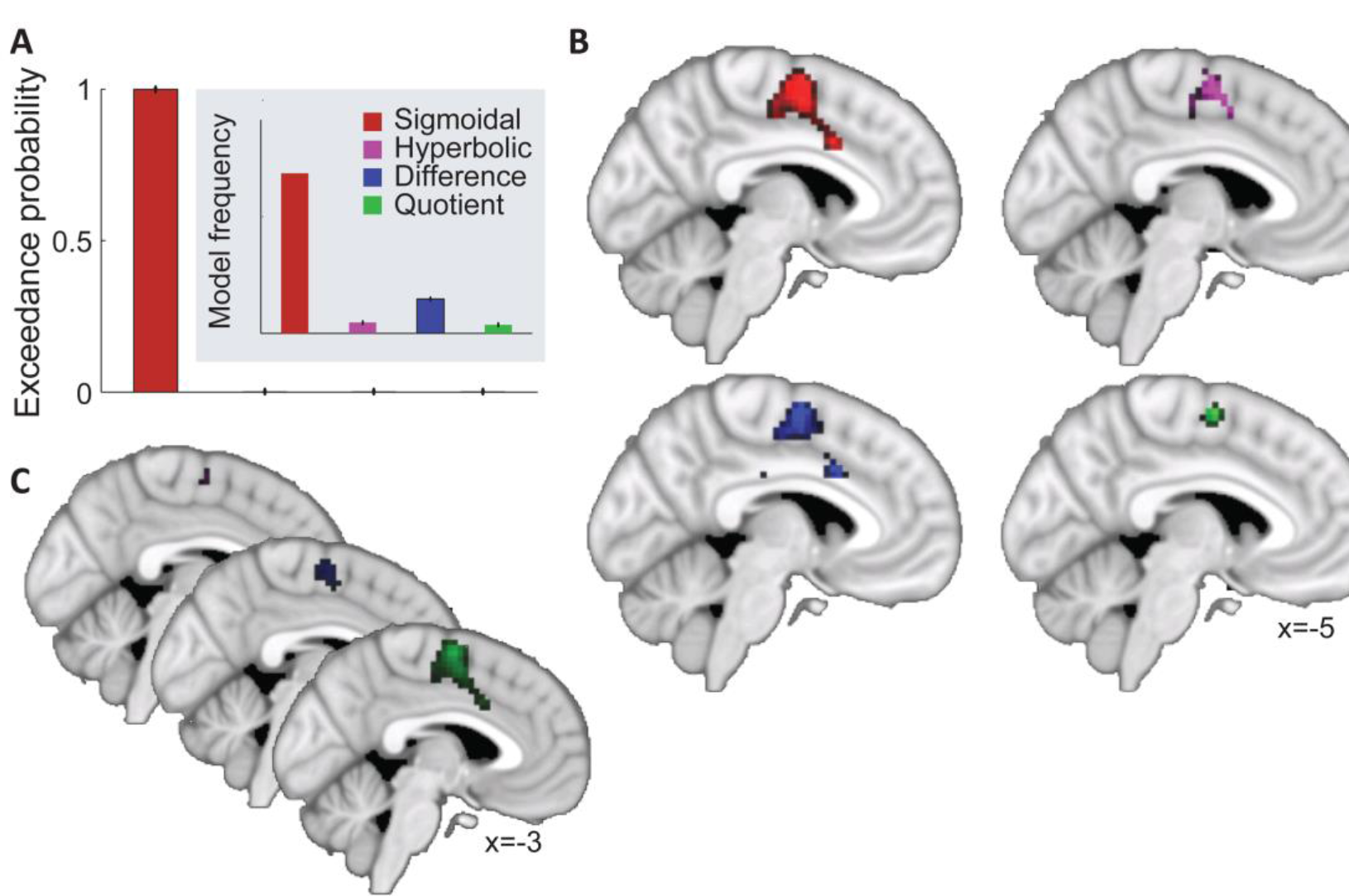
Model-derived value describes choice signals more accurately than model-freevalue. **A**, Bayesian model comparison for value modelled using the sigmoidal model, hyperbolic model and two parameter-free descriptions of value-reward minus effort,and reward divided by effort. The sigmoidal model captures choice behaviour best. **B**, Comparison of the BOLD response to value difference for the behavioural sigmoidal model (red; like **Figure 2D**), the hyperbolic model (purple), and the parameter-free descriptions of value (blue: reward-effort; green: reward/effort; all shown at p<0.001 uncorr.). The three alternative contrasts reveal a similar network albeit less strongly. **C**, Crucially, the sigmoidal model provides a significantly better description of the BOLD signal inSMA, extending into caudaldACC, compared to all other models. Purple: sigmoidal versus hyperbolic; blue: sigmoidal versus parameter-free subtraction; green: sigmoidal versus parameter-free division (shown at p<0.001 uncorr.).

### Relating neural and behavioural effects of value difference

If it was indeed the case that the regions identified to encode value difference are involved in choice computation and as a result, inform behaviour, the BOLD value signal should systematically relate to behavioural measures of choice performance (Jocham et al., 2012; Kolling et al., 2012). To test this, we used the behavioural measure of the effect of value difference, *β*_V_, as derived from the logistic regression analysis above (equation (2)). Importantly, before fitting *β*_V_, model-derived subjective values were scaled between [0,1] for all participants so that any difference in the fitted regression coefficient *β*_V_ indicated how strongly value difference influenced behavioural choices in a given participant. *β*_V_ reflects how consistently participants choose the subjectively more valuable option. In other words, this parameter captures how strongly value rather than noise determines choice behaviour. To examine whether the size of the neural value difference signal carried behavioural relevance, the behavioural weights *β*_V_ were then used as a covariate for the value difference contrast in a second level group analysis. At the whole-brain level we thus identified regions where the encoding of value difference was significantly modulated by how strongly participants' choices were driven by subjective value (**Figure 2F**). This analysis was restricted to the regions that encoded value difference at the first level. For illustration of the effect, the neural signature of value difference (regression coefficients for chosen versus unchosen value at the peak time of 6s) was plotted against *β*_V_ (**Figure 2F**).

### Reward maximization versus effort minimization

In our task, reward maximization is in conflict with effort minimization in almost all trials because the option that has a higher reward value is also associated with a higher effort level. To capture the separate behavioural influences of reward and effort for each participant, another logistic regression analysis was conducted, but now both the difference in offer magnitudes and in efforts were entered into the design matrix, rather than just their combination into value as in equation (2):

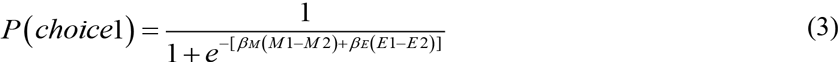

Here, *β*_M_ is the weight or precision with which reward magnitude difference (M1-M2) influences choice, *β*_E_ and is the weight (precision) with which effort difference (E1-E2) influences choice.

Next, to identify which brain regions might bias the choice computation either towards reward or away from physical effort, we performed two independent tests. First, we used the behaviourally defined weights for effort, *-β*_E_, as a covariate on the second level, to identify regions where the encoding of effort difference scales with how “effort averse” participants were. In such regions, a larger difference between chosen and unchosen effort signals would indicate that participants avoid efforts more strongly (**Figure 4B**). Based on prior work, we had a priori hypotheses about effort preferences being guided by SMA and putamen (e.g., Croxson et al., 2009; Kurniawan et al., 2010, 2013; Burke et al., 2013). Therefore, we used a small-volume correction (p<0.05) around previously established coordinates (putamen [±26, −8, −2]; SMA [4 −6 58], see (Croxson et al., 2009)). Secondly, in an analogous fashion, we used the behavioural weights for reward magnitude, *β*_M_, as a covariate on the second level to identify regions where the encoding of reward magnitude difference scales with how reward-seeking participants are. In brain regions thus identified, a larger BOLD signal difference between chosen and unchosen reward signals would imply that participants place a stronger weight on reward maximization in their choices (**Figure 4A**). Based on prior work, we expected reward magnitude comparisons to occur in vmPFC (e.g., Kable and Glimcher, 2007; Boorman et al., 2009; Philiastides et al., 2010). Therefore, we used a small-volume correction (p<0.05) around previously established coordinates [−6, 48, −8] (Boorman et al., 2009).

**Figure 4.**
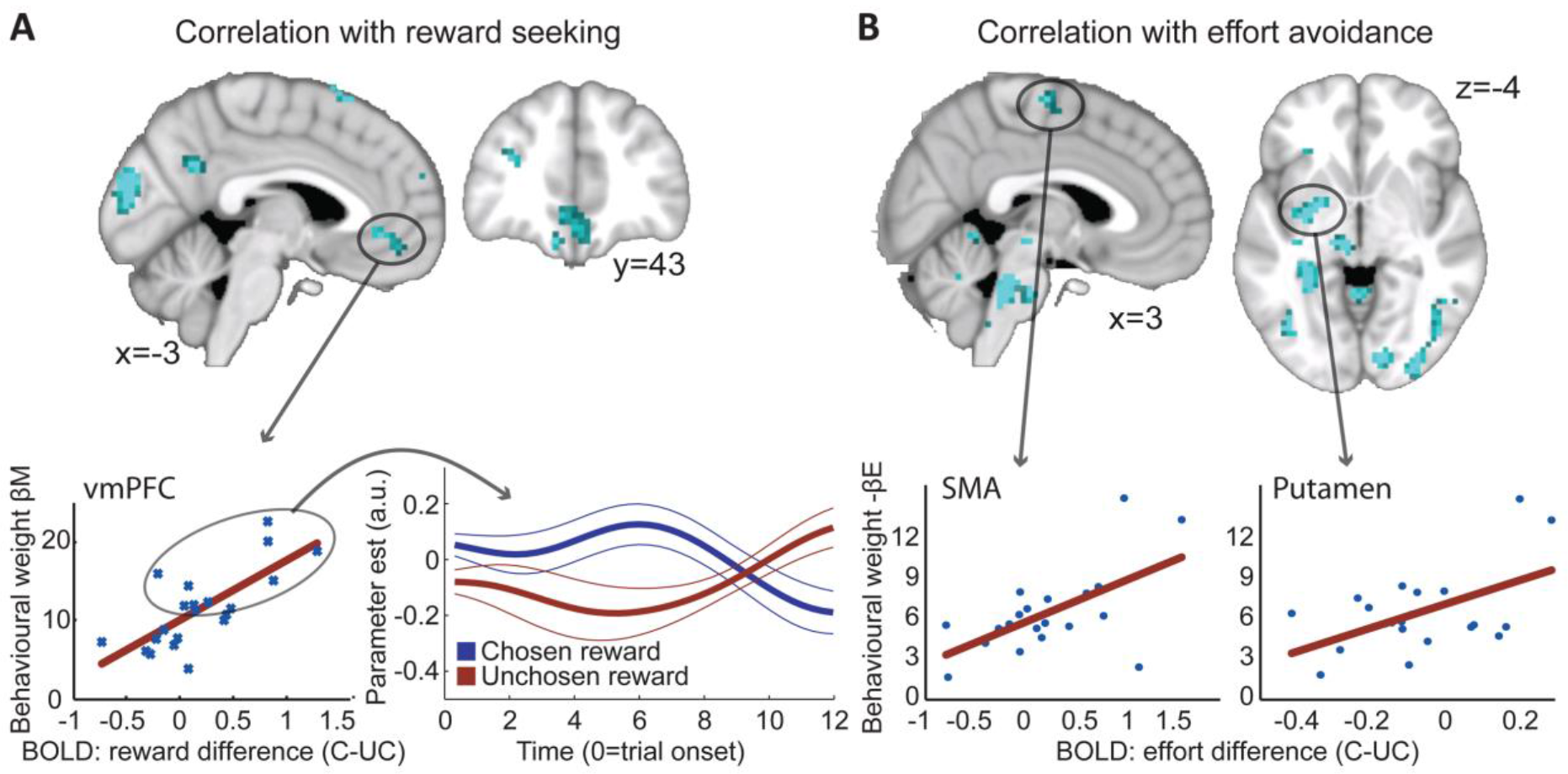
Distinct circuits bias choices towards reward maximization or effortminimization. **A**, Regions where the encoding of reward magnitude difference varied as a function of the behavioural weight participants placed on reward (β_M_; top: shown at p<0.01 uncorr.). This showed that the BOLD signal in vmPFC (SVC FWE-corr, p<0.05) reflected the difference between chosen and unchosen reward more strongly in participants who also placed a stronger weight on maximising reward (top and bottom left). While we could not identify an average reward difference coding in vmPFC across the group,the subset of participants who placed a stronger weight on reward (larger β_M_; mediansplit, ellipse) did encode the difference between the chosen and unchosen reward magnitudes (bottom right). This suggests that vmPFC might bias choices towards reward-maximization. Standard errors denote ± SEM.**B**, A very distinct network of regions including the SMA and putamen (both SVC FWE-corr, p<0.05) encoded effort difference as a function of participants' behavioural effort weight (β_E_; shown at p<0.01 uncorr.). This system was active more strongly in participants who tried to more actively avoid higher efforts and has often been associated with effort evaluation. It might counteract the vmPFC-circuit shown in A in order to achieve effort minimization,which is in constant conflict with reward maximization in our task. Correlation plots (bottom) are only shown for visual illustration of the effects for a priori regions of interest; no statistical analyses were performed on these data.

We further characterized the relationship between participants' effort sensitivity and BOLD signal changes by asking whether the neural encoding of effort difference relates to the individual distortions captured in the parameters *k* and *p* of the effort discounting function. For each trial, we compared the true effort difference between the chosen and unchosen option with the modelled subjective effort difference between the chosen and unchosen option. We took the sum of the absolute error from the best linear fit between these two variables as an index of how well our initial GLM captured subjective distortions in the evaluation of effort. We used this measure as an additional regressor for our second level analysis, in addition to *β_E_* (these two regressors are uncorrelated: r=-0.27, p=0.24). This approach had the advantage that it combined subjective effort distortions driven by both *p* and *k* into a single parameter relevant for the effort comparison (correlation of the summed errors with k: r=0.9646, p<0.001; with p: r=0.60, p=0.0043).

## Results

Human participants performed choices between options with varying rewards and physical efforts (force grips; **Figure 1A**). Our main aim was to identify areas carrying neural signatures of value comparison, which are sometimes absent on choices when all decision variables favour the same choice (Hunt et al., 2012). Therefore, for the majority of decisions, larger rewards were paired with larger efforts so that reward maximization competed with energy minimization, and the reward and effort of each option had to be combined into an integrated subjective value to derive a choice. We first tested whether both the size of reward and the associated effort of each choice option had an impact on participant's choice behaviour. A logistic regression showed that participants' choices were indeed guided by the reward magnitude and effort of both options (left reward: t_20_=-9.71, Cohen's d=-4.34, p=4.28e-08; right reward: t_20_=8.89, Cohen's d=3.98, p=1.44e-07; left effort: t_20_=7.56, Cohen's d=3.38, p=2.79e-06; right effort: t_20_=-8.37, Cohen's d=-3.74, p=2.79e-06; **Figure 1B**). As expected, larger rewards and smaller effort costs attracted choices. Overall, participants chose the higher effort option on 48 ± 2% of trials.

Next we examined whether effort was weighed up against reward; if this was the case, the influence of effort (reward) on the participant's choice would become stronger as the reward (effort) difference between the options becomes smaller. Indeed, effort had a bigger impact on choice in trials with a small compared to a large reward difference (median split; green panels in **Figure 1D**; difference in slopes: t_20_=-18.06, p=7.51e-14; small reward difference only: slope=-1.23±0.11; t_20_=-11.51, p=2.82e-10; large reward difference only: slope=0.26±0.12, t_20_=-1.95, p=0.066). The same was true for reward: its impact on choice behaviour was greater in trials with a small compared to a large effort difference (blue panels in **Figure 1D**; difference in slopes: t_20_=11.95, p=1.46e-10; small effort difference only: slope=1.65±0.05, t_20_=36.03, p=1.14e-19; large effort difference only: slope=0.38±0.12, t_20_=3.08, p=0.0059). This analysis also confirmed that effort and reward were treated as continuous variables.

Given behaviour was guided by the costs as well as the benefits associated with the two choice options we next asked whether any brain region encoded both effort and reward in a reference frame consistent with choice. Our main aim was to identify neural signatures of the choice computation: any brain region comparing the values of the two choice options should be sensitive to information about both costs and benefits. Recent work using a biophysically realistic attractor network model (Wang, 2002) suggests that the mass activity of a region computing a choice should reflect the difference of the values of both choice options (Hunt et al., 2012). In our task, a region comparing the options should hence encode (1) the inverse difference between chosen and unchosen efforts, and (2) the (positive) difference between chosen and unchosen rewards. We therefore computed the formal conjunction of these two contrasts, which is a conservative test, asking whether any region is significant in both comparisons. This test focussed on the decision phase, which was separated in time from the motor response (**Figure 1A**). We identified a cluster of activation in the SMA and in the caudal portion of dorsal anterior cingulate cortex (dACC), on the border of the anterior and posterior rostral cingulate zones (RCZa, RCZp) and area 24 (Neubert et al., 2015) (**Figure 2A**; p<0.05 cluster-level FWE-corr; peak coordinate: [-6, 11, 34], t_1,40_=4.02; SMA peak coordinate: [−9 −7 58], ti,_40_=5.29). No other regions reached FWE cluster-corrected significance (p<0.05). Notably, we did not identify any activations in the vmPFC, a region commonly identified in reward-related value computations, even at lenient statistical thresholds (p<0.01, uncorrected). Replication of this conjunction analysis in FSL, performed at the suggestion of one reviewer, obtained comparable results, with only dACC and SMA reaching cluster-level corrected significance (**Figure 2A**, green overlays). The two difference signals for effort and reward are illustrated for the BOLD time series extracted from the dACC cluster in **Figure 2B**.

These results raise the question of whether and how effort and reward are combined into an integrated value for each option-a prerequisite for testing whether any brain region encodes the comparison between subjective option values. While established models exist to examine how participants compute compound values for uncertain/risky rewards (prospect theory, (Kahneman and Tversky, 1979; Tversky and Kahneman, 1992)) and delayed rewards (hyperbolic, (Mazur, 1987; Laibson, 1997; Frederick et al., 2002)), it remains unclear how efforts and rewards are combined into a subjective measure of value. We performed several behavioural experiments to develop a behavioural model that can formally describe the range of effort discounting behaviours observed in healthy populations (Klein-Flügge et al., 2015). One key feature of this model is that it can accommodate cases when increases in effort at lower effort levels have a comparatively small effect on value, compared to increases in effort at higher effort levels (i.e., concave discounting).

When we fitted this model to the choices recorded during the scanning session, participants' behaviour was indeed best captured by an initially concave discounting shape (initially concave in 16 of 21 participants; **Figure 2C**), consistent with previous work (Klein-Flügge et al., 2015) and the intuition that effort increases are less noticeable at lower levels of effort compared to higher levels of effort.

Using the individual model fits, we then directly tested for neural signatures consistent with a value comparison between the subjective values of the two choice options. This is a slightly less conservative test than the formal conjunction of effort and reward magnitude difference described above, but we note that this test revealed a highly consistent pattern of results. We found strong evidence for a network consisting of the SMA (peak: [-9 −7 58], t_1,20_=8.64), caudal portion of dACC (peak: [−3 11 34], t_1,20_=7.1) and bilateral putamen (several peaks: left [-33 −13 4], t_1_,_20_=4.96 and [-33 −10 −2], t_1_,_20_=5.28; right [33 −1 −2], t_1_,_20_=4.96) to encode the (positive) difference in subjective value between the chosen and unchosen options (**Figure 2D**; all cluster-level FWE-corr; p<0.05). Again, comparable results were obtained using FSL (green overlays in **Figure 2D**). This network resembled regions previously described for the evaluation of physical effort, but was clearly distinct from the neural system associated with decisions about goods involving the vmPFC (Kable and Glimcher, 2007; Boorman et al., 2009; Fitzgerald et al., 2009; Philiastides et al., 2010; Hunt et al., 2012; Kolling et al., 2012; Clithero and Rangel, 2014; Strait et al., 2014).

To validate our choice of behavioural discounting model, we performed a formal model comparison and found that the sigmoidal model provided a better explanation of choice behaviour than (convex) hyperbolic discounting, previously proposed for effort discounting (Prévost et al., 2010), and two parameter-free descriptions of value ‘reward minus effort’ and ‘reward divided by effort’ (model exceedance probability: xp=1; mean of posterior distribution: mp_sigm=0.75; mp_hyp=0.05; mp_diff=0.16; mp_div=0.04; **Figure 3A**). On average, the sigmoidal model correctly predicted 88 ± 1% of choices. To examine whether our measure of value derived from the sigmoidal model also best predicted the BOLD signal, we re-calculated the value difference contrasts in an analogous way, this time modelling value using a hyperbolic or one of the two parameter-free models. The resulting whole-brain maps similarly highlighted SMA and dACC (surviving cluster-level FWE-corr., p<0.05 for the hyperbolic and difference models, n.s. for the quotient model; **Figure 3B**). But importantly, direct statistical comparison showed that the neural signal in these regions was significantly better explained by the values derived from the sigmoidal model (cluster-level FWE-corr., p<0.05 for the difference and quotient models; sigmoidal versus hyperbolic: SMA peak [-3 −7 61], t_1,19_=3.95; sigmoidal versus difference: dACC peak [-6,11,34], t_1,19_=3.28; SMA peak [-6 −7 58], t_1_,_19_=5.33; sigmoidal versus quotient: dACC peak [-6,11,34], t_1_,_19_=4.77; SMA peak [-6 −7 61], t_1,19_=6.72; **Figure 3C**). This suggests that the BOLD signal aligns with the subjective experience of effort-discounted value which was best captured using the sigmoidal model.

A crucial question is whether the observed value difference signal bears any behavioural relevance for choice, rather than potentially being a mere bi-product of a choice computation elsewhere. In the former case, one would expect that the encoding of subjective value difference relates to the strength, or ‘weight’, with which subjective value difference influenced behaviour across participants (Jocham et al., 2012; Kolling et al., 2012; Khamassi et al., 2015). Such a behavioural weight was derived for each participant using a logistic regression on the normalized model-derived subjective values. The resulting parameter estimate is the same as the inverse softmax temperature or precision and reflects how consistently participants choose the subjectively more valuable option (see Methods: in equation 2). The only region that was significant in this second-level test and also encoded value difference at the first level was the dACC (**Figure 2F**; cluster-level FWE-corr., p<0.05; peak [-3 11 31], t_1,19_=3.71). In other words, dACC encoded value difference on average across the group, and participants who exhibited a larger BOLD value difference signal in the dACC were also more consistent in choosing the subjectively better option (larger β_V_); this relationship is illustrated in **Figure 2F**.

To further probe whether the identified network of regions evaluates the choice options in a subjective manner, we examined the relationship between the subjective 'distortion’ of effort described by the parameters *k* and *p* of the individual effort discount function, and the BOLD signal related to the effort difference across participants. We calculated a measure to describe how much the true effort difference deviated from the subjectively experienced effort difference overall across trials. This 'distortion’ regressor correlated with *k* (r=0.9646, p<0.001) and *p* (r=0.60, p=0.0043), but not *β_E_*(r=-0.27, p=0.24), and was used as a second-level covariate for the effort difference contrast. GLM1 contained the efforts shown on the screen and thus should have captured the subjectively experienced effort better in participants who showed smaller effort distortions (i.e., with discounting closer to linear). Thus, in regions related to the comparison of subjective effort or effort-integrated value, we expected participants with less effort distortions to show a stronger negative effort difference signal. Indeed, we found such a positive second-level correlation with the BOLD signal in dACC and SMA, supporting the notion that effort difference is encoded in these regions in the way it subjectively influences the choice (**Figure 2G**, cluster-level FWE-corr., p<0.05; dACC peak [-3 14 31], t_1,19_=5.01, global maxima; SMA peak [-6, −7, 61], t1,19=4.08).

ACC has access to information from motor structures (Selemon and Goldman-Rakic, 1985; Dum and Strick, 1991; Morecraft and Van Hoesen, 1992, 1998; Kunishio and Haber, 1994; Beckmann et al., 2009) previously linked to evaluating effort (Croxson et al., 2009; Kurniawan et al., 2010, 2013; Burke et al., 2013), and prefrontal regions known to be involved in reward processing, such as the vmPFC and OFC (Padoa-Schioppa and Assad, 2006; Kennerley and Wallis, 2009; Levy and Glimcher, 2011; Rudebeck and Murray, 2011; Klein-Flügge et al., 2013; Chau et al., 2015; Stalnaker et al., 2015). We thus reasoned that the ACC may be a key node for the type of effort-based choice assessed in the present task. To further test this hypothesis, we sought to identify regions that mediate between reward maximisation versus effort minimization in our task.

To this end, we first extracted two separate behavioural weights reflecting participants' tendency to seek reward and avoid effort. These behavioural parameters were derived from a logistic regression with two regressors explaining how much choices were guided by the difference in reward magnitude and the difference in effort level between options (see Methods: *β*_M_ and *β*_E_ in equation 3). This is distinct from using just one regressor for the combined subjective value difference as done above (*β*_V_). Across participants, we then first identified brain regions where the encoding of chosen versus unchosen reward magnitude correlated with the weight, *β*_M_, with which choices were influenced by the difference in *reward* between the chosen and unchosen option. Secondly, we performed the equivalent test for effort, i.e. we identified regions where the neural encoding of chosen versus unchosen effort correlated with the weight,-*β*_E_, with which behaviour was guided by the difference in *effort* between the chosen and unchosen option. The two tests revealed two distinct networks of regions. First, the vmPFC encoded reward magnitude difference across subjects as a function of how much participants' choices were driven by the difference in reward between the options (SVC FWE-corr cluster-level p=0.037; peak [-6 44 −8], t_1_,_19_=2.87; **Figure 4A**). Unlike in many other tasks (Kable and Glimcher, 2007; Boorman et al., 2009; Fitzgerald et al., 2009; Philiastides et al., 2010; Hunt et al., 2012; Kolling et al., 2012; Clithero and Rangel, 2014; Strait et al., 2014), the vmPFC BOLD signal did not correlate with chosen reward or reward difference on average in the group. However, reward difference signals were on average positive for participants whose choices were more strongly driven by reward magnitudes (median split; **Figure 4A**). At the whole-brain level, the correlation of behavioural reward-weight, *β*_M_, and BOLD reward difference encoding did not reveal any activations using our FWE cluster-level corrected criterion of p<0.05. Using a lenient exploratory threshold (p=0.01, uncorr), we identified a small number of other regions including the posterior cingulate cortex (PCC) bilaterally and visual cortex (**Figure 4A**), but crucially no clusters in motor, supplementary motor or striatal regions.

By contrast, a network of motor regions including SMA and putamen encoded effort difference as a function of the individual behavioural effort weight *-β*_E_ (**Figure 4B**; SVC FWE-corr cluster-level SMA: p=0.048, peak [3 −7 58], t_1,19_=2.59; left putamen: p=0.035, peak [-27 −4 − 5], t_1_,_19_=3.39; right putamen no supra-threshold voxels). In other words, these regions encoded the difference in effort between the chosen and unchosen options more strongly in participants whose choices were negatively influenced by large effort differences, i.e. participants who were more sensitive to effort costs. Using a whole-brain FWE cluster-level-corrected threshold (p<0.05), no regions were detected in this contrast. At an exploratory threshold (p=0.01, uncorr), this contrast also highlighted regions in the brainstem, primary motor cortex, thalamus and dorsal striatum (**Figure 4B**), and thus regions previously implicated in evaluating motor costs and in recruiting resources in anticipation of effort (Croxson et al., 2009; Burke et al., 2013; Kurniawan et al., 2013), but clearly distinct from the vmPFC/PCC network identified in the equivalent test for reward above.

Taken together, our data thus show that two distinct networks centred on vmPFC versus SMA/putamen encode the reward versus effort difference as a function of how much these variables influence the final choice. Yet only the caudal portion of dACC encodes the difference in overall subjective value as a function of how much overall value influences choice. This region in dACC could therefore be a potential mediator between reward maximization and effort minimization which appear to occur in separate neural circuits.

### Functionally distinct sub-regions of medial PFC

For completeness, we also tested if any areas encode an opposite value difference signal (i.e., the inverse of the conjunction analysis and of the subjective value difference contrast performed above), reflecting the evidence *against* the chosen option and thus one notion of decision difficulty. This did not reveal any regions at our conservative (cluster-level FWE-corrected) threshold in either test. At a more lenient exploratory threshold (p=0.01 uncorr), a single common cluster in medial PFC (pre-SMA/area 9) was identified (**Figure 5**), in agreement with previous reports of negative value difference signals in this region (Wunderlich et al., 2009; Hare et al., 2011). Importantly, the location of this activation was clearly distinct from the caudal dACC region found to encode a positive value difference (**Figure 2**). Here, by contrast, value difference signals did not correlate with the strength with which subjective value difference influenced behaviour across participants (*β*_V_; no supra-threshold voxels at p=0.01 uncorr.), suggesting this region's functions during choice are separate from those that bias behaviour.

**Figure 5.**
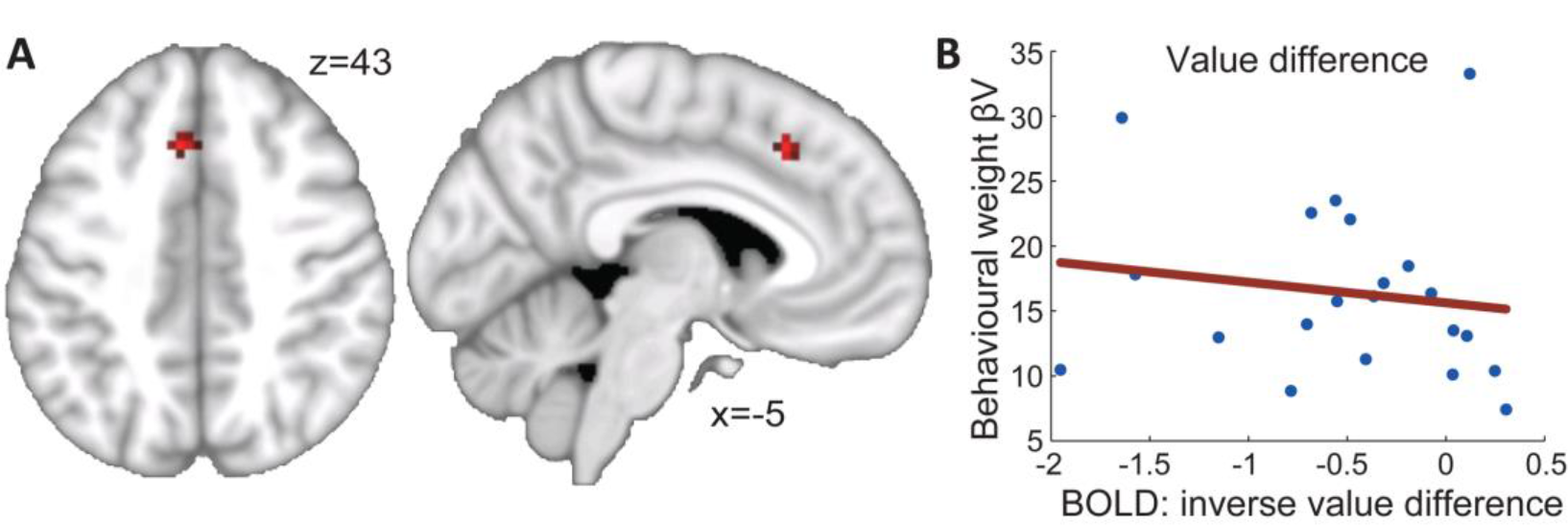
Opposite coding of relative choice value in dorsal medial frontal cortex. **A**, Regions where the BOLD signal encodes an inverse rather than a positive difference between chosen and unchosen reward magnitudes and a positive rather than an inverse difference between chosen and unchosen effort (i.e. the exact inverse of the conjunction shown in Figure 2A). The only region detected at a lenient threshold (p=0.01 uncorr.; no regions survive FWE correction) is a nearby but anatomically distinct region in medial prefrontal cortex (mPFC) previously suggested to serve as a choice comparator (Wunderlich et al., 2009; Hare et al., 2011). **B**, However, in this region, the BOLD signal does not relate to behaviour as was the case for the caudal portion of dACC (see **Figure 2F**).

## Discussion

Choices requiring the consideration of motor costs are ubiquitous in everyday life. Unlike other types of choices they require knowledge of the current state of the body and its available energy resources, to weight physical costs against potential benefits. How this trade-off might be implemented neurally remains largely unknown.

Here, we identified a region in the caudal part of dorsal anterior cingulate cortex (dACC) as the key brain region that carried the requisite signatures for effort-based choice: dACC represented the costs and benefits of the chosen relative to the alternative option, integrated effort and reward into a combined subjective value signal, computed the subjective value difference between the chosen relative to the alternative option, and activity here correlated with the degree to which participants' choices were driven by value.

### ACC integrates effort and reward information

Work from several lines of research suggests ACC may be a key region for performing cost-benefit integration for effort-based choice. For example, lesions to ACC (but not OFC) result in fewer choices of a high effort/high reward compared to a low effort/low reward option: yet such animals still choose larger reward options when effort costs for both options are equated, implying ACC is not essential when decisions can be solved only by reward (Walton et al., 2003, 2009; Schweimer and Hauber, 2005; Rudebeck et al., 2006; Floresco and Ghods-Sharifi, 2007). BOLD responses in human ACC reflect the integrated value of effort-based options in the absence of choice (Croxson et al., 2009). Further, single neuron recordings from ACC encode information about both effort and reward (Shidara and Richmond, 2002; Kennerley et al., 2009, 2011), and integrate costs and benefits into a value signal (Hillman and Bilkey, 2010; Hosokawa et al., 2013; Hunt et al., 2015). ACC thus appears to have a critical role in integrating effort and reward information to derive the subjective value of performing a particular action.

### ACC encodes a choice comparison signal

However, from the aforementioned work it remained unclear whether cost-benefit values of different choice options are actually *compared* in ACC, or whether reward and effort may be compared in separate neural structures and the competition resolved between areas. When one choice option is kept constant, the value of the changing option correlates perfectly with the value difference between the options (Kurniawan et al., 2010; Prévost et al., 2010; Bonnelle et al., 2016), which precludes distinguishing between valuation and value comparison processes. This is similarly true when only one option is offered and accepted/rejected (Bonnelle et al., 2016). We here varied both options' values from trial to trial, which allowed us to identify a choice *comparison* signal in the ACC, and thus the essential neural signature implicating this area in decision making. Firstly we show a region in the caudal portion of dACC encodes separate difference signals for effort and reward. The direction of these difference signals aligns with their respective effect on value, with effort decreasing and reward increasing an option's overall value. Secondly, we demonstrate a comparison signal between integrated option values. We used a novel behavioural model (Klein-Flügge et al., 2015) to characterize participants' individual tendency to discount reward given the level of motor costs. Using the resultant model-derived subjective values, we identified the dACC as a region encoding a combined value difference signal. Indeed, our model provided a better characterization of the BOLD signal than other models of effort discounting, and dACC activity was related to individuals' 'distortions' of effort. This resolves an important question showing that effort and reward information are indeed brought together within a single region to inform choice.

Finally, this value comparison signal also varied as a function of how much value influenced choices across participants. This result further strengthens the idea that the dACC plays a crucial role in guiding choice, rather than merely representing effort or reward information. In our task, no other region exhibited similar dynamics even at lenient thresholds.

### Influences from ‘effort’ and ‘reward’ circuits

Nevertheless, an important question remains: do the regions that preferentially encode /reward or effort have any influence on choice? To examine this question, we looked for regions that explain participants' tendency to avoid effort, or to seek reward. This analysis revealed two distinct circuits. Whereas signals in vmPFC reflected the relative benefits as a function of how reward-driven participants' choices were, a network more commonly linked to action selection and effort evaluation (Croxson et al., 2009; Kurniawan et al., 2010, 2013; Prévost et al., 2010; Burke et al., 2013; Bonnelle et al., 2016)-including SMA and putamen-encoded relative effort as a function of how much participants tried to avoid energy expenditure. It will be of interest to examine in future work how these circuits interact, and how different modulatory systems contribute to this interplay (see e.g., Varazzani et al., 2015). This question should be extended to situations when different costs coincide or different strategies compete (see Burke et al., 2013 for one recent example), or when information about effort and reward has to be learnt (Skvortsova et al., 2014; Scholl et al., 2015).

### Converging evidence for multiple decision circuits

Our results contribute to an emerging literature demonstrating the existence of multiple decision systems in the brain which are flexibly recruited based on the type of decision (Rushworth et al., 2012). One well-studied system concerns choices where costs are directly tied to outcomes (e.g., risk, delay). During this type of choice, vmPFC encodes the difference between the chosen and unchosen options' cost-benefit value (Kable and Glimcher, 2007; Boorman et al., 2009; Philiastides et al., 2010; Hunt et al., 2012; Kolling et al., 2012), consistent with the decision impairments observed after vmPFC lesions (Noonan et al., 2010; Camille et al., 2011a, 2011c). Other types of choices, however, rely on other networks (Kolling et al., 2012; Hunt et al., 2014; Wan et al., 2015). In the present study, decisions required the integration of motor costs and we show that for this dACC, rather than vmPFC, plays a more central role. VmPFC did not encode overall value or the difference in value between the options in our task; in our hands, vmPFC evidenced no information about effort costs, consistent with previous proposals (Prévost et al., 2010; Skvortsova et al., 2014).

### Functionally dissociable anatomical sub-regions of mPFC

The location in the dACC identified here is distinct from a more anterior and dorsal region in medial frontal cortex (in or near pre-SMA) where BOLD encodes the opposite signal: a negative value difference (Wunderlich et al., 2009; Hare et al., 2011). It is also more posterior than a dACC region involved in foraging choices (Kolling et al., 2012). The cluster of activation identified here extends from the cingulate gyrus dorsally into the lower bank of the cingulate sulcus, and it is sometimes also referred to it as midcingulate cortex (MCC; Procyk et al., 2016) or rostral cingulate zone (Ridderinkhof et al., 2004). According to a recent connectivity-based parcellation, our activation is on the border of areas RCZa (34%), RCZp (33%) and area 24 (48%) (Neubert et al., 2015). While it shares some voxels with the motor cingulate regions in humans (Amiez and Petrides, 2014), most parts of our cluster are more ventral and located in the gyral portion of ACC (see also Kolling et al., 2016 for a discussion of functionally dissociable activations in ACC).

### Relevance for disorders of motivation

Our findings in the dACC speak to an important line of research showing deficits in effort-based decision making in a number of disorders including depression, negative symptom schizophrenia and apathy (Levy and Dubois, 2006; Cléry-Melin et al., 2011; Treadway et al., 2012, 2015; Fervaha et al., 2013; Gold et al., 2013; Hartmann et al., 2014; Pizzagalli, 2014; Yang et al., 2014; Bonnelle et al., 2015). Patients with these disorders often show a reduced ability to initiate effortful actions to obtain reward. Crucially, they also exhibit abnormalities in ACC and basal-ganglia circuits, as well as other regions processing information about the autonomic state, including the amygdala and some brainstem structures (Drevets et al., 1997; Botteron et al., 2002; Levy and Dubois, 2006). Furthermore, individuals with greater behavioural apathy scores show enhanced recruitment of precisely the circuits implicated in the present study, including SMA and cingulate cortex, when deciding to initiate effortful behaviour (Bonnelle et al., 2016). This is interesting because apathy correlates with increased effort sensitivity (*β_E;_* Bonnelle et al., 2016), and we found that individuals with increased effort sensitivity showed enhanced recruitment of SMA and brainstem regions for encoding the effort difference (Fig 3B). In other words, when committing to a larger (relative) effort, these circuits were more active in people who were more sensitive to effort. As discussed in Bonnelle et al. (2016), we cannot infer cause and effect but it is possible that the neural balance between activations in reward and effort systems might be different in individuals with greater sensitivity to efforts (such as apathetic individuals). This may be why these people avoid choosing effortful options more often than others. It also provides a possible connection between the network's specific role in effort-based choice and its functional contribution to everyday life behaviours.

## Acknowledgements

MCKF, SWK and KF were supported by the Wellcome Trust (MCKF: 086120/Z/08/Z; SWK 096689/Z/11/Z; KF: 088130/Z/09/Z and 091593/Z/10/Z); SB was supported by the European Research Council (ERC; ActSelectContext 260424). Conceived and designed the experiments: MCK SWK SB. Performed the experiments: MCK. Analysed the data: MCK. Contributed reagents/materials/analysis tools: KF. Wrote the paper: MCK SWK KF SB. We would like to thank Tim Behrens, Laurence Hunt, Matthew Rushworth, Marco Wittmann and all our lab members for helpful discussions on the data, and the imaging and IT teams at the FIL and Sobell Department for their support with data acquisition and handling.

## References

Amiez C, Petrides M (2014) Neuroimaging evidence of the anatomo-functional organization of the human cingulate motor areas. Cereb Cortex N Y N 1991 24:563–578.

Andersson JL, Hutton C, Ashburner J, Turner R, Friston K (2001) Modeling geometric deformations in EPI time series. NeuroImage 13:903–919.

Beckmann M, Johansen-Berg H, Rushworth MFS (2009) Connectivity-Based Parcellation of Human Cingulate Cortex and Its Relation to Functional Specialization. J Neurosci 29:1175–1190.

Birn RM, Diamond JB, Smith MA, Bandettini PA (2006) Separating respiratory-variation-related fluctuations from neuronal-activity-related fluctuations in fMRI. NeuroImage 31:1536–1548.

Bonnelle V, Manohar S, Behrens T, Husain M (2016) Individual Differences in Premotor Brain Systems Underlie Behavioral Apathy. Cereb Cortex N Y NY 26:807–819.

Bonnelle V, Veromann K-R, Burnett Heyes S, Lo Sterzo E, Manohar S, Husain M (2015) Characterization of reward and effort mechanisms in apathy. J Physiol Paris 109:16–26.

Boorman ED, Behrens TEJ, Woolrich MW, Rushworth MFS (2009) How green is the grass on the other side? Frontopolar cortex and the evidence in favor of alternative courses of action. Neuron 62:733–743.

Botteron KN, Raichle ME, Drevets WC, Heath AC,Todd RD (2002) Volumetric reduction in left subgenual prefrontal cortex in early onset depression. Biol Psychiatry 51:342–344.

Burke CJ, Brünger C, Kahnt T, Park SQ, Tobler PN (2013) Neural Integration of Risk and Effort Costs by the Frontal Pole: Only upon Request. J Neurosci Off J Soc Neurosci 33:1706–1713.

Camille N, Griffiths CA, Vo K, Fellows LK, Kable JW (2011a) Ventromedial Frontal Lobe Damage Disrupts Value Maximization in Humans. J Neurosci 31:7527–7532.

Camille N, Tsuchida A, Fellows LK (2011b) Double dissociation of stimulus-value and action-value learning in humans with orbitofrontal or anterior cingulate cortex damage. J Neurosci Off J Soc Neurosci 31:15048–15052.

Camille N, Tsuchida A, Fellows LK (2011c) Double dissociation of stimulus-value and action-value learning in humans with orbitofrontal or anterior cingulate cortex damage. J Neurosci Off J Soc Neurosci 31:15048–15052.

Chau BKH, Sallet J, Papageorgiou GK, Noonan MP, Bell AH, Walton ME, Rushworth MFS (2015) Contrasting Roles for Orbitofrontal Cortex and Amygdala in Credit Assignment and Learning in Macaques. Neuron 87:1106–1118.

Cléry-Melin M-L, Schmidt L, Lafargue G, Baup N, Fossati P, Pessiglione M (2011) Why don't you try harder? An investigation of effort production in major depression. PloS One 6:e23178.

Clithero JA, Rangel A (2014) Informatic parcellation of the network involved in the computation of subjective value. Soc Cogn Affect Neurosci 9:1289–1302.

Croxson PL, Walton ME, O'Reilly JX, Behrens TEJ, Rushworth MFS (2009) Effort-based cost-benefit valuation and the human brain. J Neurosci Off J Soc Neurosci 29:4531–4541.

Drevets WC, Price JL, Simpson JR Jr, Todd RD, Reich T, Vannier M, Raichle ME (1997) Subgenual prefrontal cortex abnormalities in mood disorders. Nature 386:824–827.

Dum RP, Strick PL (1991) The origin of corticospinal projections from the premotor areas in the frontal lobe. J Neurosci Off J Soc Neurosci 11:667–689.

Eklund A, Nichols T, Knutsson H (2015) Can parametric statistical methods be trusted for fMRI based group studies? ArXiv151101863 Math Stat Available at:http://arxiv.org/abs/1511.01863 [Accessed May 26, 2016].

Fervaha G, Foussias G, Agid O, Remington G (2013) Neural substrates underlying effort computation in schizophrenia. Neurosci Biobehav Rev 37:2649–2665.

Fitzgerald THB, Seymour B, Dolan RJ (2009) The role of human orbitofrontal cortex in value comparison for incommensurable objects. J Neurosci Off J Soc Neurosci 29:8388–8395.

Floresco SB, Ghods-Sharifi S (2007) Amygdala-prefrontal cortical circuitry regulates effort-based decision making. Cereb Cortex N Y N 1991 17:251–260.

Frederick S, Loewenstein G, O'Donoghue T (2002) Time Discounting and Time Preference: A Critical Review. J Econ Lit 40:351–401.

Friston K, Mattout J, Trujillo-Barreto N, Ashburner J, Penny W (2007) Variational free energy and the Laplace approximation. NeuroImage 34:220–234.

Glover GH, Li T-Q, Ress D (2000) Image-based method for retrospective correction of physiological motion effects in fMRI: RETROICOR. Magn Reson Med 44:162–167.

Gold JM, Strauss GP, Waltz JA, Robinson BM, Brown JK, Frank MJ (2013) Negative Symptoms of Schizophrenia Are Associated with Abnormal Effort-Cost Computations. Biol Psychiatry.

Hare TA, Schultz W, Camerer CF, O'Doherty JP, Rangel A (2011) Transformation of stimulus value signals into motor commands during simple choice. Proc Natl Acad Sci U S A 108:18120–18125.

Hartmann MN, Hager OM, Reimann AV, Chumbley JR, Kirschner M, Seifritz E, Tobler PN, Kaiser S (2014) Apathy But Not Diminished Expression in Schizophrenia Is Associated With Discounting of Monetary Rewards by Physical Effort. Schizophr Bull.

Hayden BY, Platt ML (2010) Neurons in anterior cingulate cortex multiplex information about reward and action. J Neurosci Off J Soc Neurosci 30:3339–3346.

Hillman KL, Bilkey DK (2010) Neurons in the rat anterior cingulate cortex dynamically encode cost-benefit in a spatial decision-making task. J Neurosci Off J Soc Neurosci 30:7705–7713.

Hosokawa T, Kennerley SW, Sloan J, Wallis JD (2013) Single-neuron mechanisms underlying cost-benefit analysis in frontal cortex. J Neurosci Off J Soc Neurosci 33:17385–17397.

Hunt LT, Behrens TE, Hosokawa T, Wallis JD, Kennerley SW (2015) Capturing the temporal evolution of choice across prefrontal cortex. eLife:e11945.

Hunt LT, Dolan RJ, Behrens TEJ (2014) Hierarchical competitions subserving multi-attribute choice. Nat Neurosci 17:1613–1622.

Hunt LT, Kolling N, Soltani A, Woolrich MW, Rushworth MFS, Behrens TEJ (2012) Mechanisms underlying cortical activity during value-guided choice. Nat Neurosci 15:470–476.

Hutton C, Josephs O, Stadler J, Featherstone E, Reid A, Speck O, Bernarding J, Weiskopf N (2011) The impact of physiological noise correction on fMRI at 7T. NeuroImage 57:101–112.

Jocham G, Hunt LT, Near J, Behrens TEJ (2012) A mechanism for value-guided choice based on the excitation-inhibition balance in prefrontal cortex. Nat Neurosci 15:960–961.

Kable JW, Glimcher PW (2007) The neural correlates of subjective value during intertemporal choice. Nat Neurosci 10:1625–1633.

Kahneman D, Tversky A (1979) Prospect theory-analysis of decision under risk. Econometrica 47:263–291.

Kennerley SW, Behrens TEJ, Wallis JD (2011) Double dissociation of value computations in orbitofrontal and anterior cingulate neurons. Nat Neurosci 14:1581–1589.

Kennerley SW, Dahmubed AF, Lara AH, Wallis JD (2009) Neurons in the frontal lobe encode the value of multiple decision variables. J Cogn Neurosci 21:1162–1178.

Kennerley SW, Wallis JD (2009) Evaluating choices by single neurons in the frontal lobe: outcome value encoded across multiple decision variables. Eur J Neurosci 29:2061–2073.

Kennerley SW, Walton ME, Behrens TEJ, Buckley MJ, Rushworth MFS (2006) Optimal decision making and the anterior cingulate cortex. Nat Neurosci 9:940–947.

Khamassi M, Quilodran R, Enel P, Dominey PF, Procyk E (2015) Behavioral Regulation and the Modulation of Information Coding in the Lateral Prefrontal and Cingulate Cortex. Cereb Cortex N Y N 1991 25:3197–3218.

Klein-Flügge MC, Barron HC, Brodersen KH, Dolan RJ, Behrens TEJ (2013) Segregated Encoding of Reward-Identity and Stimulus-Reward Associations in Human Orbitofrontal Cortex. J Neurosci Off J Soc Neurosci 33:3202–3211.

Klein-Flügge MC, Kennerley SW, Saraiva AC, Penny WD, Bestmann S (2015) Behavioral modeling of human choices reveals dissociable effects of physical effort and temporal delay on reward devaluation. PLoS Comput Biol 11:e1004116.

Kolling N, Behrens T, Wittmann M, Rushworth M (2016) Multiple signals in anterior cingulate cortex. Curr Opin Neurobiol 37:36–43.

Kolling N, Behrens TEJ, Mars RB, Rushworth MFS (2012) Neural Mechanisms of Foraging. Science 336:95–98.

Kunishio K, Haber SN (1994) Primate cingulostriatal projection: Limbic striatal versus sensorimotor striatal input. J Comp Neurol 350:337–356.

Kurniawan IT, Guitart-Masip M, Dayan P, Dolan RJ (2013) Effort and valuation in the brain: the effects of anticipation and execution. J Neurosci Off J Soc Neurosci 33:6160–6169.

Kurniawan IT, Seymour B, Talmi D, Yoshida W, Chater N, Dolan RJ (2010) Choosing to make an effort: the role of striatum in signaling physical effort of a chosen action. J Neurophysiol 104:313–321.

Laibson D (1997) Golden Eggs and Hyperbolic Discounting. Q J Econ 112:443–478.

Levy DJ, Glimcher PW (2011) Comparing apples and oranges: using reward-specific and reward-general subjective value representation in the brain. J Neurosci Off J Soc Neurosci 31:14693–14707.

Levy R, Dubois B (2006) Apathy and the Functional Anatomy of the Prefrontal Cortex-Basal Ganglia Circuits. Cereb Cortex 16:916–928.

Luk C-H, Wallis JD (2009) Dynamic encoding of responses and outcomes by neurons in medial prefrontal cortex. J Neurosci Off J Soc Neurosci 29:7526–7539.

Matsumoto K, Suzuki W, Tanaka K (2003) Neuronal correlates of goal-based motor selection in the prefrontal cortex. Science 301:229–232.

Mazur J (1987) An adjusting procedure for studying delayed reinforcement. In: Quantitative analyses of behavior: Vol. 5. The effect of delay and of intervening events on re-inforcement value (Commons M, Mazur J, Nevin J, Rachlin H, eds). Erlbaum.

Morecraft RJ, Van Hoesen GW (1992) Cingulate input to the primary and supplementary motor cortices in the rhesus monkey: evidence for somatotopy in areas 24c and 23c. J Comp Neurol 322:471–489.

Morecraft RJ, Van Hoesen GW (1998) Convergence of limbic input to the cingulate motor cortex in the rhesus monkey. Brain Res Bull 45:209–232.

Neubert F-X, Mars RB, Sallet J, Rushworth MFS (2015) Connectivity reveals relationship of brain areas for reward-guided learning and decision making in human and monkey frontal cortex. Proc Natl Acad Sci U S A 112:E2695–2704.

Noonan MP, Walton ME, Behrens TEJ, Sallet J, Buckley MJ, Rushworth MFS (2010) Separate value comparison and learning mechanisms in macaque medial and lateral orbitofrontal cortex. Proc Natl Acad Sci 107:20547–20552.

Padoa-Schioppa C, Assad JA (2006) Neurons in the orbitofrontal cortex encode economic value. Nature 441:223–226.

Pastor-Bernier A, Cisek P (2011) Neural correlates of biased competition in premotor cortex. J Neurosci Off J Soc Neurosci 31:7083–7088.

Penny W, Kiebel S, Friston K (2003) Variational Bayesian inference for fMRI time series. NeuroImage 19:727–741.

Philiastides MG, Biele G, Heekeren HR (2010) A mechanistic account of value computation in the human brain. Proc Natl Acad Sci U S A 107:9430–9435.

Pizzagalli DA (2014) Depression, stress, and anhedonia: toward a synthesis and integrated model. Annu Rev Clin Psychol 10:393–423.

Prévost C, Pessiglione M, Météreau E, Cléry-Melin M-L, Dreher J-C (2010) Separate valuation subsystems for delay and effort decision costs. J Neurosci Off J Soc Neurosci 30:14080–14090.

Procyk E, Wilson CRE, Stoll FM, Faraut MCM, Petrides M, Amiez C (2016) Midcingulate Motor Map and Feedback Detection: Converging Data from Humans and Monkeys. Cereb Cortex N Y N 1991 26:467–476.

Rangel A, Hare T (2010) Neural computations associated with goal-directed choice. Curr Opin Neurobiol 20:262–270.

Ridderinkhof KR, Ullsperger M, Crone EA, Nieuwenhuis S (2004) The role of the medial frontal cortex in cognitive control. Science 306:443–447.

Rudebeck PH, Behrens TE, Kennerley SW, Baxter MG, Buckley MJ, Walton ME, Rushworth MFS (2008) Frontal cortex subregions play distinct roles in choices between actions and stimuli. J Neurosci Off J Soc Neurosci 28:13775–13785.

Rudebeck PH, Murray EA (2011) Balkanizing the primate orbitofrontal cortex: distinct subregions for comparing and contrasting values. Ann N Y Acad Sci 1239:1–13.

Rudebeck PH, Walton ME, Smyth AN, Bannerman DM, Rushworth MFS (2006) Separate neural pathways process different decision costs. Nat Neurosci 9:1161–1168.

Rushworth MF, Kolling N, Sallet J, Mars RB (2012) Valuation and decision-making in frontal cortex: one or many serial or parallel systems? Curr Opin Neurobiol Available at: http://www.ncbi.nlm.nih.gov/pubmed/22572389 [Accessed May 27, 2012].

Scholl J, Kolling N, Nelissen N, Wittmann MK, Harmer CJ, Rushworth MFS (2015) The Good, the Bad, and the Irrelevant: Neural Mechanisms of Learning Real and Hypothetical Rewards and Effort. J Neurosci Off J Soc Neurosci 35:11233–11251.

Schweimer J, Hauber W (2005) Involvement of the rat anterior cingulate cortex in control of instrumental responses guided by reward expectancy. Learn Mem Cold Spring Harb N 12:334–342.

Selemon LD, Goldman-Rakic PS (1985) Longitudinal topography and interdigitation of corticostriatal projections in the rhesus monkey. J Neurosci 5:776–794.

Shidara M, Richmond BJ (2002) Anterior cingulate: single neuronal signals related to degree of reward expectancy. Science 296:1709–1711.

Skvortsova V, Palminteri S, Pessiglione M (2014) Learning to minimize efforts versus maximizing rewards: computational principles and neural correlates. J Neurosci Off J Soc Neurosci 34:15621–15630.

Stalnaker TA, Cooch NK, Schoenbaum G (2015) What the orbitofrontal cortex does not do. Nat Neurosci 18:620–627.

Strait CE, Blanchard TC, Hayden BY (2014) Reward value comparison via mutual inhibition in ventromedial prefrontal cortex. Neuron 82:1357–1366.

Treadway MT, Bossaller NA, Shelton RC, Zald DH (2012) Effort-based decision-making in major depressive disorder: a translational model of motivational anhedonia. J Abnorm Psychol 121:553–558.

Treadway MT, Peterman JS, Zald DH, Park S (2015) Impaired effort allocation in patients with schizophrenia. Schizophr Res 161:382–385.

Tversky A, Kahneman D (1992) Advances in Prospect Theory: Cumulative Representation of Uncertainty. J Risk Uncertain 5:297–323.

Varazzani C, San-Galli A, Gilardeau S, Bouret S (2015) Noradrenaline and dopamine neurons in the reward/effort trade-off: a direct electrophysiological comparison in behaving monkeys. J Neurosci Off J Soc Neurosci 35:7866–7877.

Walton ME, Bannerman DM, Alterescu K,Rushworth MFS (2003) Functional specialization within medial frontal cortex of the anterior cingulate for evaluating effort-related decisions. J Neurosci Off J Soc Neurosci 23:6475–6479.

Walton ME, Groves J, Jennings KA, Croxson PL, Sharp T, Rushworth MFS, Bannerman DM (2009) Comparing the role of the anterior cingulate cortex and 6-hydroxydopamine nucleus accumbens lesions on operant effort-based decision making. Eur J Neurosci 29:1678–1691.

Walton ME, Kennerley SW, Bannerman DM, Phillips PEM, Rushworth MFS (2006) Weighing up the benefits of work: behavioral and neural analyses of effort-related decision making. Neural Netw Off J Int Neural Netw Soc 19:1302–1314.

Wan X, Cheng K, Tanaka K (2015) Neural encoding of opposing strategy values in anterior and posterior cingulate cortex. Nat Neurosci 18:752–759.

Wang X-J (2002) Probabilistic decision making by slow reverberation in cortical circuits. Neuron 36:955–968.

Ward NS, Frackowiak RSJ (2003) Age-related changes in the neural correlates of motor performance. Brain 126:873–888.

Weiskopf N, Hutton C, Josephs O, Deichmann R (2006) Optimal EPI parameters for reduction of susceptibility-induced BOLD sensitivity losses: a whole-brain analysis at 3 T and 1.5 T. NeuroImage 33:493–504.

Wunderlich K, Rangel A, O'Doherty JP (2009) Neural computations underlying action-based decision making in the human brain. Proc Natl Acad Sci U S A 106:17199–17204.

Yang X-H, Huang J, Zhu C-Y, Wang Y-F, Cheung EFC, Chan RCK, Xie G-R (2014) Motivational deficits in effort-based decision making in individuals with subsyndromal depression, first-episode and remitted depression patients. Psychiatry Res 220:874–882.

